# Spike-independent infection of human coronavirus 229E in bat cells

**DOI:** 10.1101/2021.09.18.460924

**Authors:** Marcus G Mah, Martin Linster, Dolyce HW Low, Zhuang Yan, Jayanthi Jayakumar, Firdaus Samsudin, Foong Ying Wong, Peter J Bond, Ian H Mendenhall, Yvonne CF Su, Gavin JD Smith

**Affiliations:** Programme in Emerging Infectious Diseases, Duke-NUS Medical School, Singapore; Bioinformatics Institute, Agency for Science, Technology, and Research, Singapore; Department of Biological Sciences, National University of Singapore, Singapore; Centre for Outbreak Preparedness, Duke-NUS Medical School, Singapore; SingHealth Duke-NUS Global Health Institute, SingHealth Duke-NUS Academic Medical Centre, Singapore; Duke Global Health Institute, Duke University, Durham, NC 27710, USA

**Keywords:** evolution, pandemic, receptor usage, zoonotic, spike

## Abstract

Bats are the reservoir for numerous human pathogens including coronaviruses. The factors leading to the emergence and sustained transmission of coronaviruses in humans are poorly understood. An outstanding question is how coronaviruses can accomplish a host switch with a likely mismatch between the surface protein spike of a bat virus and the human cellular receptor at the time of zoonotic virus transmission. To identify potential novel evolutionary pathways for zoonotic virus emergence, we serially passaged six human 229E isolates in a newly established *Rhinolophus lepidus* (horseshoe bat) kidney cells and analyzed viral genetic changes. Here we observed extensive deletions within the spike and ORF4 genes of five 229E viruses after passaging in bat cells. As a result, spike protein expression and infectivity of human cells was lost in 5 of 6 viruses but the capability to infect bat cells was maintained. Only viruses that expressed the spike protein could be neutralized by 229E spike-specific antibodies in human cells, whereas there was no neutralizing effect on viruses that do not express the spike protein inoculated on bat cells. However, one isolate acquired an early stop codon abrogating spike expression but maintaining infection in bat cells. Upon passaging this isolate in human cells, spike expression was restored due to acquisition of nucleotide insertions amongst virus subpopulations. Spike-independent infection of coronaviruses provides an alternative mechanism for viral maintenance in bats that does not rely on the compatibility of viral surface proteins and cellular entry receptors.

## Introduction

Coronaviruses (CoVs) are positive-sense RNA viruses with large genomes ranging from 27kb to 32kb. CoVs are members of the *Coronaviridae* family, within the *Orthocoronavirinae* subfamily and are divided into four genera, *Alphacoronavirus, Betacoronavirus, Deltacoronavirus*, and *Gammacoronavirus* (1). Four seasonal CoVs circulate globally, 229E, NL63, OC43, and HKU1 (2). Together they account for 10% to 30% of upper respiratory tract infections in adults (3). Recently, three zoonotic coronaviruses have emerged in human populations - SARS-CoV-1 (4,5), MERS-CoV (6), and SARS-CoV-2 (7). Based on genetic analyses, 229E, NL63, SARS-CoV-1, SARS-CoV-2, and MERS-CoV have ancestral origins in bats (8,9). Recently, 229E-related bat coronaviruses have been isolated from *Hipposideros* and *Rhinolophus* bats across Africa (10,11). Temporal analyses of 229E sequences from bats and humans have estimated that 229E viruses diverged from bat ancestral viruses more than 130 years ago (12). The detection of 229E-related viruses in captive alpaca and dromedary suggests that 229E might have evolved from bats to humans via camelids as intermediate hosts (13,14).

Bats harbor a variety of pathogenic viruses including Ebola virus, Nipah virus, Hendra virus, lyssaviruses and coronaviruses without displaying discernable symptoms of infection (15,16). In order to infect humans, bat-derived CoVs need to overcome host-species barriers and balance conservation and novel acquisition of genes while maintaining infectivity, replication, and spread (17). Entry of CoVs into host cells is generally mediated by the binding of the spike protein to host receptors which in humans comprise angiotensin-converting enzyme 2 (ACE2) for SARS-CoV-1 (18), SARS-CoV-2 (19), and NL63 (20); dipeptidyl peptidase-4 (DPP4) for MERS-CoV (21); and aminopeptidase N for 229E (22). It is unclear how bat CoVs can cross between hosts without a match between the surface protein spike and the cellular receptor that is involved in binding and internalization of virions. We here show evidence for the replication of 229E CoVs that have lost spike gene expression during serial passaging in a kidney cell line derived from *Rhinolophus lepidus* bats (Blyth’s horseshoe bat). These viruses remain capable of infecting bat cells but are unable to infect human cells.

## Results

To identify possible adaptive mutations, five clinical isolates of 229E coronavirus obtained from nasopharyngeal swabs and one reference strain (VR740) were cultured in human colon adenocarcinoma (Caco2) cells twice (C2 virus). These viruses were then serially passage ten times (C2R1 to C2R10) in a newly established *Rhinolophus lepidus* kidney (Rhileki) cell line (Fig S1). During virus passaging in Rhileki cells, an increase in viral genome copies from inoculum to 6 days post inoculation (dpi) was observed except for isolate SG/1197/2010 (Fig 1A). The inoculation of Caco2 and Rhileki cells with C2 viruses resulted in infection as detected by nucleocapsid immunofluorescence staining (Figure S2) and titration in the respective cell line (Fig 1B and 1C). However, C2R10 viruses were only able to infect Rhileki cells but not Caco2 cells, except for isolate SG/2613/2011. Viral titers of C2 viruses in Caco2 cells ranged from 6.8×10^4^ to 3.2×10^7^ TCID_50_/ml (Fig 1B), whereas C2R1 virus titers in Rhileki cells ranged from 1×10^2^ to 2.68×10^4^ TCID_50_/ml and C2R10 viruses displayed virus titers 1×10^2^ to 1.58×10^4^ TCID_50_/ml (Fig 1C). Interestingly, when C2R10 viruses were inoculated in Caco2 cells (C2R10C1), only isolate SG/2613/2011 produced a virus titer of 1.8×10^6^ TCID_50_/ml (Fig 1B). As an additional proof of viral genomic replication, double-stranded RNA was detected at 1 dpi in a Caco2 and Rhileki cells (Fig S3).

**Fig 1.**
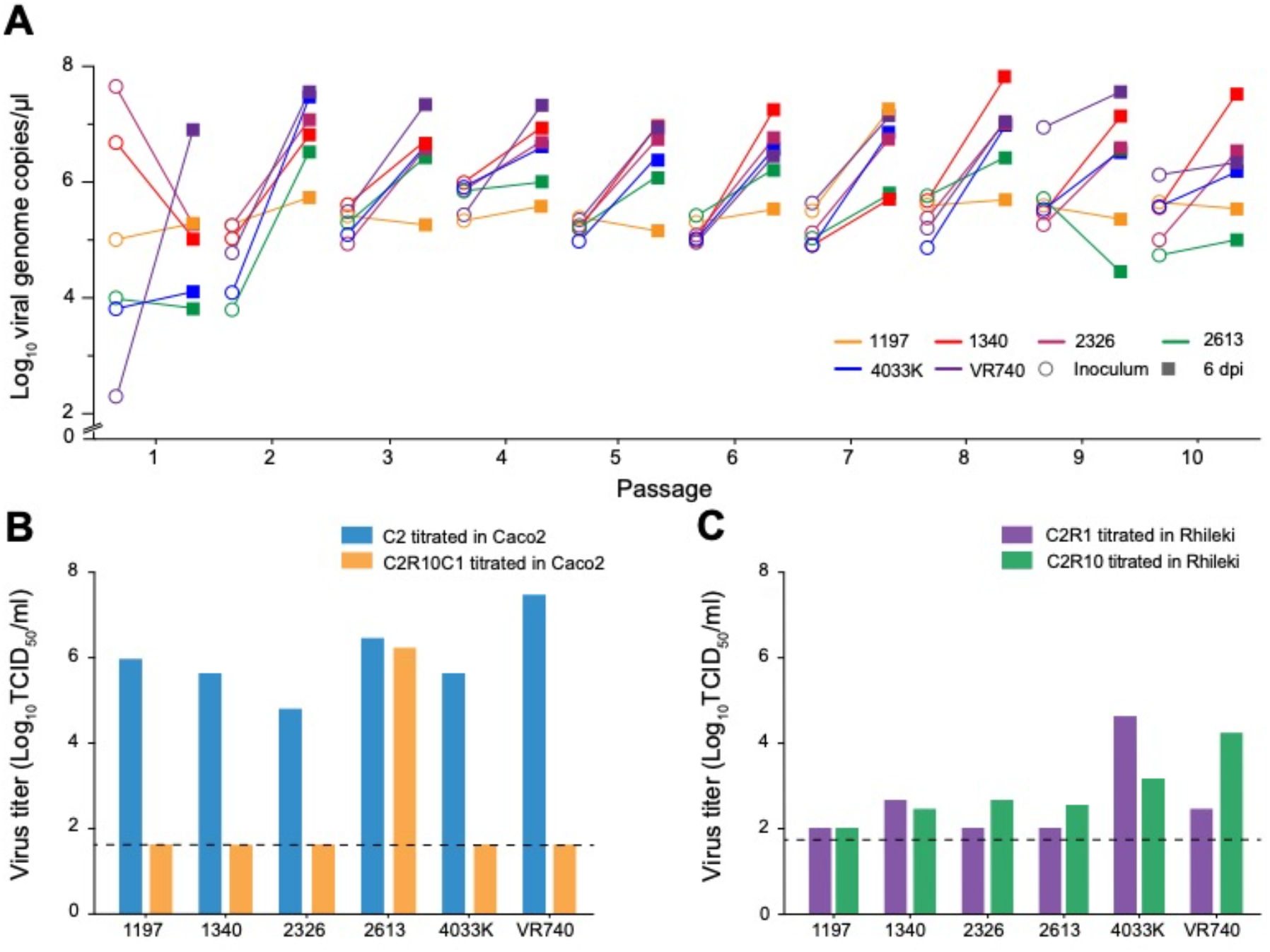
Rhileki cells are susceptible and permissive to 229E infection. (A) Serial passage of six 229E isolates at MOI = 0.01 in Rhileki cells. Viral genome copies of the inoculum (circles) and at 6 days post inoculation (square) are plotted. The inoculum for subsequent passages was standardized based on genome copy numbers. (B) Viral titers (TCID_50_/ml) in Caco2 cells of C2, C2R10 and C2R10C1 viruses after serial passage. (C) Viral titers (TCID_50_/ml) in Rhileki cells of C2R1 and C2R10 viruses after serial passage. Black dashed lines represent the limit of detection.

To determine the genetic changes that arose during passaging in Rhileki cells, we performed next-generation sequencing of C2R1, C2R5, and C2R10 viruses for the six 229E isolates (Fig. 2 and Fig. S4). Surprisingly, deletions in the spike and ORF4 gene region were observed for SG/1197/2010 (2878 nt) and TZ/4033K/2017 (449 nt) during passage one. For SG/1340/2011 (443 nt) and SG/2326/2011 (302 nt and 22 nt), deletions were detected in the C2R5 virus. The reference strain UK/VR740/1973 displayed a deletion of 2461 nt only during or before passage ten. No deletions were observed for SG/2613/2011. All deletions were confirmed by Sanger sequencing with isolate-specific flanking primers (Table S1) and minor variants were assessed by assembling reads that map to the deletions in the consensus sequences (Table S2). Potential viral subpopulations were detected in 0.65% to 2.64% of all reads. Between passage one and five, the deletion in SG/1197/2010 enlarged from 2878 nt to 3656 nt and remained constant upon further passaging. Similarly, deletions in SG/1340/2011 extended from 443 nt to 3407 nt between C2R5 and C2R10 viruses. While an additional deletion of 663 nt was found in isolate TZ/4033/2017 during or before passage five which enlarged to 2704 nt during or before passage ten, the initial deletion of 449 nt grew to 544 nt during or before passage five. Notably, the receptor binding domains (RBD) in the spike were deleted in SG/1197/2010, SG/1340/2010, TZ/4033K/2017, and UK/VR740/1973. Although no deletions were observed for SG/2613/2011, there were nucleotide insertions at position 21,069 that resulted in a premature stop codon at positions 21,097–21,099. Strikingly, as described above, the observed deletions and early stop codon did not prevent infection of Rhileki cells during the serial passage, indicating that virus infection was occurring independent of the spike protein.

**Fig 2.**
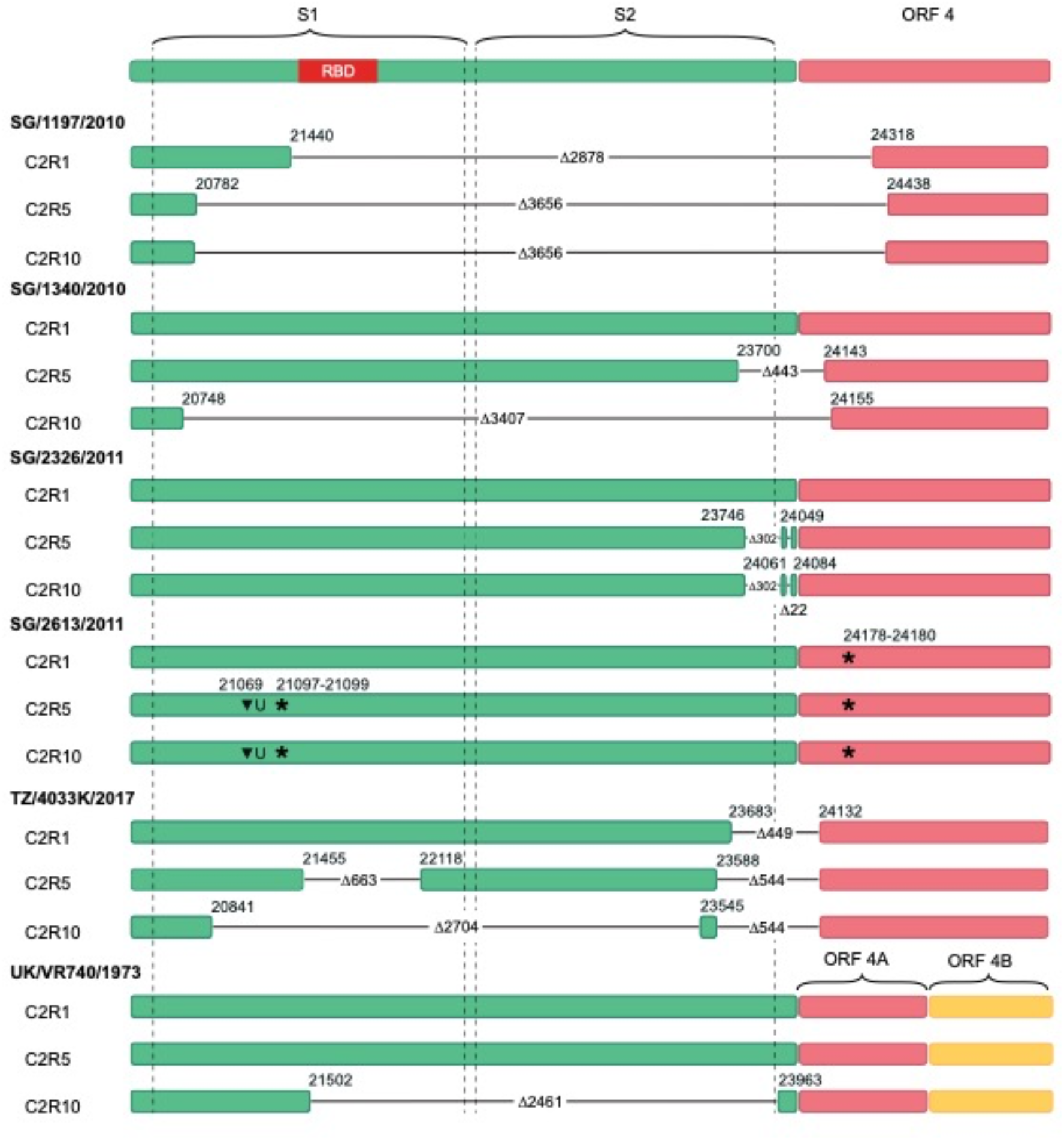
Schematic of nucleotide sequences within the spike and ORF 4 gene regions. Colored filled rectangles represent parts of the genome in C2R1, C2R5, and C2R10 viruses that are present in the respective passage. Virus isolates names are in bold. Horizontal black lines indicate deleted parts of the genome with the respective size denoted by Δ, starts, and ends of the deletions are indicated by the respective nucleotide position.

To verify the expression of spike protein in Rhileki cells, we generated a post-immunization polyclonal rabbit serum directed against the spike protein of 229E virus UK/VR740/1973 and determined the relative expression of spike protein of the serially passaged 229E isolates in Western blot analyses. No spike protein was detected in C2R1, C2R5, and C2R10 viruses, except for SG/2326/2011 C2R5 (Fig 3A). Low levels of nucleocapsid protein were observed, particularly in C2R1 viruses, corresponding to the relatively lower virus titers observed in Rhileki cells. To test if continued viral growth in Rhileki cells was because of virus subpopulations (minor variants) that encode for the spike protein, we performed virus neutralization tests (VNT) using our 229E anti-spike polyclonal serum. Full neutralization was achieved at 10μg/ml for isolates SG/1340/2010 and SG/2326/2011, 20μg/ml for isolates SG/2613/2011 and TZ/4033K/2017, 40μg/ml for SG/1197/2010, and 80μg/ml for UK/VR740/1973 for C2 viruses in Caco2 cells (Fig 3B). The six C2R10 viruses were able to infect Rhileki cells regardless of pre-incubation with spike-specific antibodies at average titers of 2.0×10^2^ TCID_50_/ml for SG/1197/2010, 3.2×10^3^ TCID_50_/ml for SG/2326/2011, 1.0×10^4^, 7.6×10^3^, 6.8×10^3^ TCID_50_/ml for SG/2613/2011, TZ/4033K/2017, UK/VR740/1973, respectively, and 6.4×10^5^ TCID_50_/ml for SG/1340/2010 at 3 days post infection. The sole C2R10C1 virus SG/2613/2011 that was able to infect Caco2 cells was neutralized at 20μg/ml in line with its C2 counterpart.

**Fig 3.**
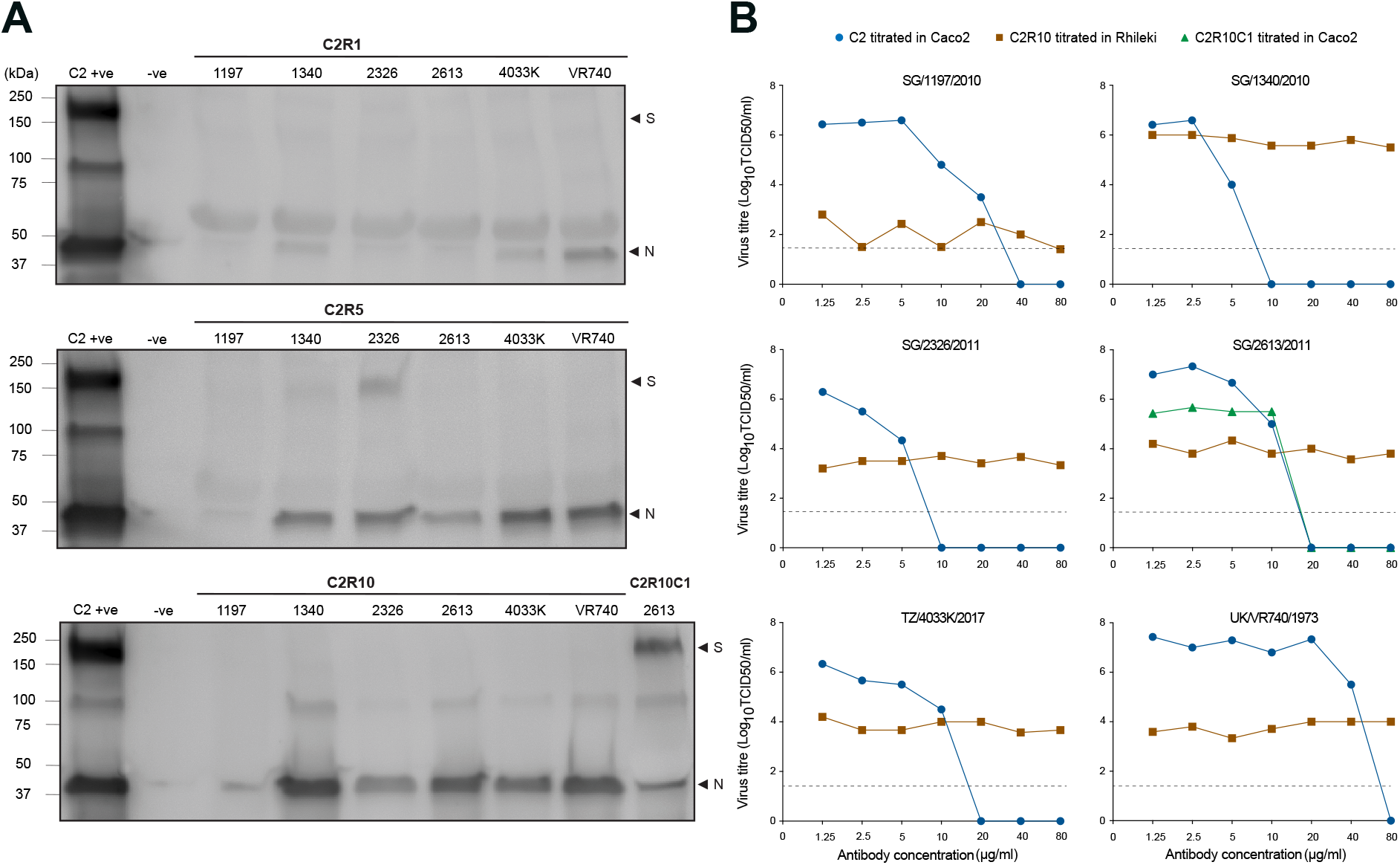
Spike protein expression and virus neutralization. (A) Western blot images of culture supernatants of Rhileki cells and Caco2 cells infected with C2R1, C2R5, C2R10, and C2R10C1, respectively, and stained with 229E anti-spike (S) and anti-nucleocapsid protein (N) antibodies. (B) Virus neutralization test, virus titers (TCID_50_/ml) of C2 viruses in Caco2 (blue), C2R10 viruses in Rhileki (brown), C2R10C1 virus in Caco2 (green). 229E anti-spike rabbit polyclonal sera at concentrations 1.25μg/ml to 80μg/ml were incubated with the respective virus 1 hour before inoculation. Blacked dashed lines indicate the limit of detection.

To test if the spike and ORF4 deletions affect sub-genomic messenger RNA (sgmRNA) transcription, we probed the unique leader-body junction (LBJ) sequences of sgmRNA 2 and 4 (Table S1) from extracted total RNA of C2 and C2R10 virus infected Caco2 and Rhileki cells, respectively (Fig S5A). While the expected PCR product size of 162 nucleotides (sgmRNA 2) and 507 nucleotides (sgmRNA 4) were obtained from C2 virus infected Caco2 cells, there was variation in band sizes in C2R10 viruses that correspond to the varying sized deletions detected in the viral genomic RNA (Fig S5B). Sanger sequencing confirmed the presence of three types of LBJs, canonical sgmRNA 2, 4, and a non-canonical LBJ was detected in SG/2326/2011 at nucleotides 23,502–23,558 of the spike open-reading frame before the putative start codon for protein X.

Given the absence of spike protein detection in our previous assays, we visualized the remaining amino acids to the 3D structures of the spike and ORF4 proteins using analogues from SARS-CoV-2 to probe whether the remaining sequences in C2R10 viruses could form a functional open reading frame (Fig S6A). Only isolate SG/2326/2011 retained an intact spike ectodomain. However, the lack of transmembrane (TM) and heptad repeat 2 (HR2) regions that are involved in membrane fusion may impair its function. The ORF4 protein is a viroporin (23). The TM domain of ORF4 is present in all but isolate SG/1197/2010 (Fig S6B). Furthermore, exogeneous treatment of Rhileki cells with *Vibrio cholerae* neuraminidase that results in the removal of surface glycans prior to infection with C2R10 viruses did not result in a consistent reduction in virus titers (data not shown).

To investigate the potential effect of mutations elsewhere in the viral genome, we identified single nucleotide polymorphisms (SNPs) in various passages (Table S3). Most SNPs (10/16) occurred in the non-structural proteins 2, 3, 4, 8, 10, and 15. There were fifteen SNPs that resulted in a non-synonymous mutation, of which four were fixed as the major variant at passage one, nine at passage five and all fifteen at passage ten. Interestingly, isolates SG/1340/2010 and SG/2326/2011 acquired mutation A4854G (D621G in NSP3) independently, whereas SG/2326/2011 and TZ/4033K/2017 obtained C25177T (L54F in the membrane protein). In addition, SG/2613/2011 acquired mutation C24200T that resulted in an early stop codon (Q30* in ORF4). Further investigation is required to determine if these mutations have any biological effect.

Only the C2R10 virus of isolate SG/2613/2011 resulted in infection of Caco2 cells (Fig. 1B) and the resulting C2R10C1 virus was positive for spike expression in Western blot analysis (Fig 3A). Therefore, we examined the sequencing reads for SG/2613/2011 spike gene, which did not display any deletions during virus passaging. Instead, virus subpopulations with insertions of one to three nucleotides at position 21,069 that resulted in loss and reinstatement of the spike open reading frame were observed (Fig 4). The C2R5 virus population had three variants, the wild type (GNFY–NE), the majority variant (GNFL– Q*) with a one nucleotide insertion resulting in a stop codon at positions 21,097–21,099, and a final variant (GNFFIAL*) with a two nucleotide insertion and a stop codon at position 21,078–21,080. The wild type was not detected in C2R10, possibly due to low sequencing coverage (Fig S4). In the C2R10C1 virus of SG/2613/2011, both the wild type (GNFY–NE) and a new variant (GNFFY–NE) with a three nucleotide insertion that reinstates the spike open reading frame was present, reiterating that spike is essential for efficient virus infection of human cells.

**Fig 4.**
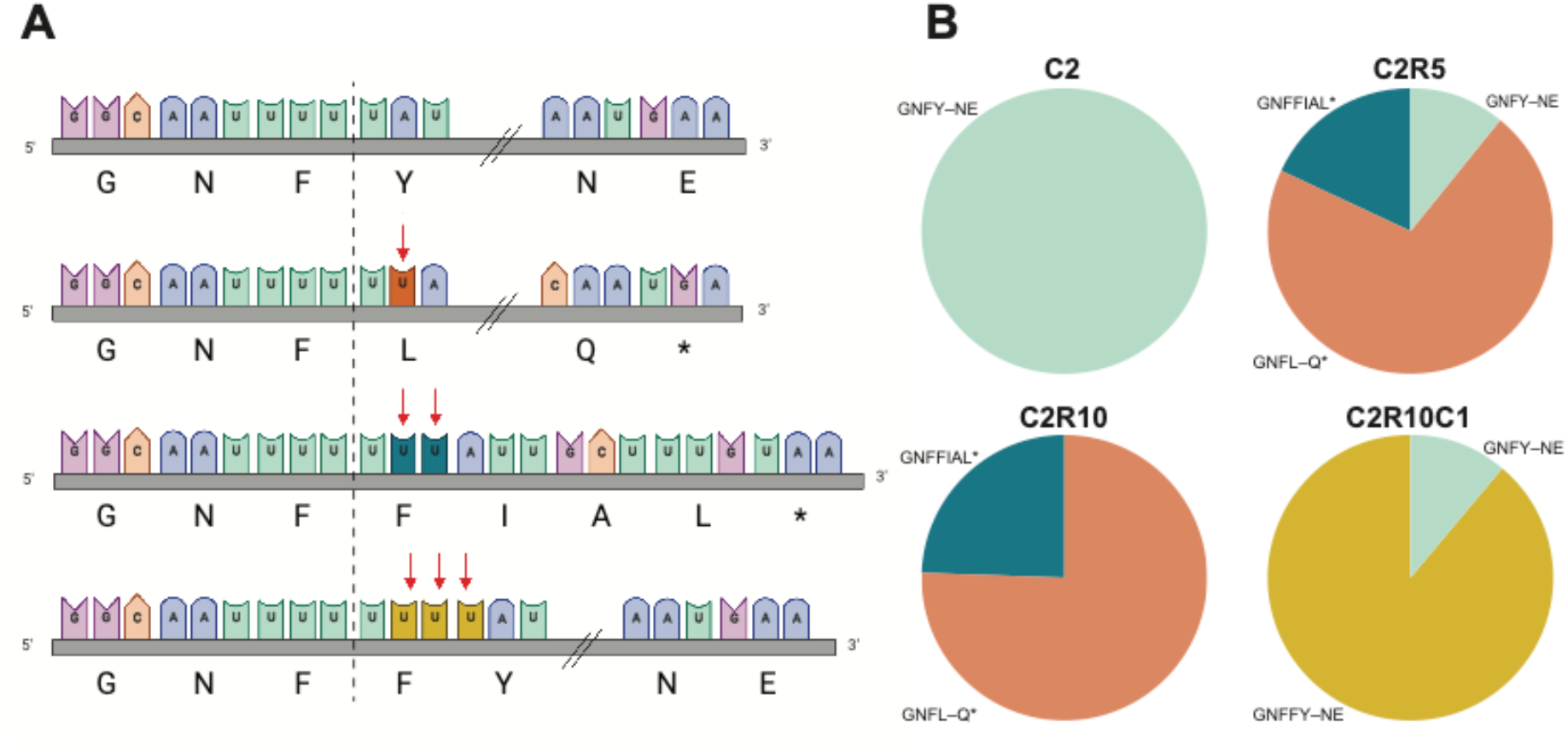
Genetic diversity of SG/2613/2011 spike gene across passages. (A) Nucleotide sequences from position 21,059–21,100 and the corresponding amino acid translation within the S1 domain of the spike gene. Depending on the passage of the virus, the diversity is comprised of one to three uracil insertions at nucleotide position 21,064 that alter the reading frame compared to the wildtype sequence (turquoise, line 1), GNFY–NE. With one (orange, line 2) or two (dark green, line 3) uracil insertions the reading frame shifts and leads to an early stop codon GNFL–Q* (E177*) and GNFFIAL* (L171*), respectively. At three (mustard, line 4) uracil insertions, the reading frame is restored with an additional phenylalanine GNFFY–NE. Double forward slash (//) indicates 8 amino acid residues not shown for clarity of presentation. (B) Pie charts describe the percentage of viral genome reads with various uracil insertions for C2 (100% no insertions, turquoise, n = 8,268), C2R5 (11.1% no insertions; 70.9% single insertion, orange; 18.0% double insertion, dark green, n = 323) C2R10 (75.8% single insertion; 24.2% double insertion, n = 33), and C2R10C1 (11.2% no insertions; 88.8% triple insertion, mustard, n = 347) viruses. Figure was created in BioRender.com.

## Discussion

Two independent mechanisms for cell entry by coronaviruses are known, receptor dependent cathepsin-mediated endocytosis and transmembrane serine protease direct cell entry (24). Both mechanisms rely on a matching viral spike protein and cellular host receptor. Human CoVs can engage a variety of human cellular receptors, e.g. SARS-CoV-2 can bind to human ACE2 but only a subset of SARS-CoV-2 related bat coronaviruses, including WIV1 and RaTG13, recognize human ACE2 (25,26). In addition, the spike protein of some bat coronaviruses including isolates RaTG13 and RmYN02 were shown to be unable to bind cognate bat ACE2 receptors (27,28), suggesting that compatibility between spike and receptor may not be necessary for virus maintenance in bats. The ability to bind human ACE2 has been mapped to residues within the RBD that are deleted or mutated in the majority of bat coronaviruses (26,29–31). Furthermore, receptor-independent entry of coronaviruses has been described for murine hepatitis virus (32), MERS-CoV (33), and SARS-CoV-2 (34). Recently, MERS-like bat coronaviruses have been described that do not recognize DPP4 but engage ACE2 (35). Therefore, the question arises of how these bat coronaviruses infect cells that do not express a compatible receptor.

Deletions in the spike and ORF4 regions occurred as early as the first passage upon serial passaging of human coronavirus 229E in Rhileki cells. Critically, the expression of the spike protein was absent in all C2R10 viruses despite sustained replication in Rhileki cells. This intriguing finding strongly suggests an alternative spike-independent mode of entry. Virus titers of C2R10 viruses were consistently lower in Rhileki cells compared to C2 viruses in Caco2 cells, suggesting that high replicative fitness of CoVs in bats might not necessarily be an evolutionary advantage (36) and is not required for virus maintenance in bat populations.

We propose spike-independent fusion of viral and cellular membranes in bat cells that is facilitated by the removal of spike protein expression that act as a spacer and sterically hinder membranes to spontaneously fuse, which has been recently observed for extracellular vesicles (37). In addition, macropinocytosis can be induced by CoV infection, can lead to productive virus replication (38), and is known as a mechanism to internalize large vesicles (39). This more generic mechanism might help to explain general features of infection observed in bats such as low levels of viraemia (40) and a dampened immune response even to highly pathogenic viruses (41). Alphacoronaviruses can infect a wider range of bats compared to betacoronaviruses (42) and spike-independent infection might offer the required flexibility for a wider host range.

This study reports the abrogation of spike protein expression consistently among six human 229E isolates in a bat kidney cell line. Although this phenomenon has not been observed in nature it might represent a novel evolutionary pathway for zoonotic virus spillover. It is plausible that small deletions or individual mutations have the functionally same effect and lead to a mismatch of spike and receptor, since viruses with large deletions in the spike gene have not been isolated so far. Consequently, the extensive deletions observed in this study might be due to the additional selective pressure for a relative smaller genome size that is more pronounced in our serial passaging cell culture experiment compared to the repeated infection of an organism in nature. The alternative mechanism to introduce an early stop codon by inserting one or two uracil nucleotides in isolate SG/2613/2011 is evidence that the absence of the spike protein, rather than the actual modification on the nucleotide level, confers an important evolutionary advantage in bat cells. Interestingly, the insertion of a third uracil restored the reading frame to synthesize spike protein with an additional amino acid residue when passaged on human cells, suggesting the importance of spike expression for infection of human cells. However, the infection dynamics in Rhileki cells likely favored replication of a shorter viral genome in the absence of other selection pressures present in a natural setting.

While the order Chiroptera contains various bat species that together harbour an astounding diversity of coronaviruses, it is unclear if spike-independent infection is restricted to Rhileki cells, *Rhinolophus lepidus*, horseshoe bats, other bat species, is organ-specific e.g. kidney or includes other groups of animals, such as rodents. Furthermore, it is not clear if spike-independent infection is restricted to 229E, (alpha-) coronaviruses, bat-borne viruses or presents an even more generic way of asymptomatic infection of a larger group of viruses. It is also possible that a viral protein other than spike binds to cell surface glycans in bats. Finally, it is yet to be determined which exact cellular mechanism allows for the internalization of 229E virions that do not contain spike proteins on their surface.

## Methods

### Establishment of the Rhileki cell line

Kidney tissue of a *Rhinolophus lepidus* bat (NUS-IACUC B01/12) was processed into cellular mixtures by mechanical disruption through a 100 μm cell strainer. One-hundred thousand cells were plated in a T-25 Corning CellBIND flask containing maintenance media (RPMI with L-glutamine, 10% FBS, 2.5% HEPES, 1x antibiotics-antimycotics) at 37 °C, 5% CO_2_ for expansion. Cells were then serially diluted in 384-well plates. Wells containing single cell patches were selected, trypsinized and serially diluted additional two times before expansion. A spontaneously immortalized clonal cell line was selected and named Rhileki for *<u>Rhi</u>nolophus <u>le</u>pidus* <u>ki</u>dney which is currently maintained until passage 30.

### Virus culture and serial passage

Isolates SG/1197/2010, SG/1340/2010, SG/2326/2011, SG/2613/2011 were obtained from a study in Singapore (43). Isolate TZ/4033K/2017 was obtained from a study conducted in Tanzania. UK/VR740/1973 was procured from the American Type Culture Collection (ATCC). All viruses were cultured for 3 days in Caco2 cells (ATCC, HTB-37) at 33°C in Dulbecco’s Modified Eagle Medium GlutaMAX containing 3% fetal bovine serum (FBS) and 1% penicillin-streptomycin. Viruses were titrated for 5 days in a 96-well format, plates were fixed with 100% methanol, blocked with 3% bovine serum albumin (BSA), stained with 1E7 primary monoclonal antibody (Eurofins Ingenasa, Spain) binding to the 229E nucleocapsid protein and a goat anti-mouse secondary antibody conjugated with fluorescein isothiocyanate (Abcam, United Kingdom). TCID50 titers were calculated according to the method of Reed and Muench (44).

Rhileki cells were seeded at a density of 2 x 10^6^ in T25 flasks. The virus inoculum at MOI 0.01 was incubated at 33°C for 2 hours. The inoculum was removed, cells washed once with phosphate buffered saline (PBS) and medium was replenished (DMEM, 1% sodium pyruvate, 1% non-essential amino acids, and 0·5% BSA). Supernatant was collected at time points 0, 3, 6, 9 dpi. At 9 dpi, the cells were scraped, pelleted, and stored in −80°C. For serial passages, viral genome copies in the supernatant of 6 dpi were adjusted to contain 10^5^ to 10^6^ genome equivalents in the subsequent inoculum. Caco2 and Rhileki cells were seeded into two-well chambered coverglass on the day before infection. Caco2 and Rhileki cells were infected with C2 or C2R10 viruses respectively for 2 hours at 33°C. Cells were fixed and permeabilized with 4% paraformaldehyde and 0.1% Triton X-100 at 1 dpi, and stained with J2 antibody (Abcam, United Kingdom) that detects for double-stranded RNA.

### RNA extraction and PCR

Briefly, total RNA was extracted from 150 μl of supernatant using the Direct-zol RNA MiniPrep kit (Zymo, USA) according to the manufacturer’s instructions. Samples were analyzed by real-time quantitative reverse-transcription-PCR using primers specific for 229E nucleocapsid. Nucleocapsid sequences were cloned and cycle threshold values were converted into viral genome copies per microliter based on regression analysis of plasmid dilutions.

### Whole-genome sequencing

Complementary DNA libraries of passage one, five, and ten were constructed using TruSeq RNA Library Prep Kit (Illumina, USA) according to the manufacturer’s instruction. The quality of the libraries was verified using the RNA bioanalyzer (Agilent, USA) and normalized using the KAPA Library Prep Kit (Roche, Switzerland). The libraries were sequenced using a 250 bp flow cell on the Illumina MiSeq System. Raw NGS reads were trimmed by Trimmomatic v0.39 to remove adaptors and low-quality reads (45). The sequence of each C2 virus isolate was used as reference to assemble trimmed and raw reads derived from Rhileki passage. The assembly was done using the BWA-MEM algorithm in UGENE v34 (46). Coverage, depth and statistics of the assembled reads were visualized in Geneious R9.1.8 and Integrative Genomics Viewer (47). Consensus sequences were aligned using MAFFT v7.222 (48) and annotated in Geneious R9.1.8.

### Genetic analysis

Trimmed and untrimmed reads were compared to flag deleted regions in the genome. To confirm the deletions, flanking primers were designed (Table S1). Using isolate specific primers, cDNA synthesis was performed using Superscript III First-Strand Synthesis System (ThermoFisher, USA). This was followed by PCR, a 50μl mixture consisted of cDNA, sense/anti-sense primers (10μM), 10x reaction buffer, Pfu polymerase (Promega, USA), and 10mM dNTP mix (ThermoFisher, USA). The PCR was conducted at 95 °C for 2 mins, 35 cycles at 95°C for 1 min, 47°C for 30 secs, 72°C for 2 mins, and final extension at 72°C for 5 mins. PCR products were visualized using a 1.5% gel and sequences confirmed by Sanger sequencing (Macrogen, South Korea).

### Sub-genomic messenger RNA Sanger sequencing

As part of the replication cycle of CoVs, translation of structural proteins is preceded by discontinuous transcription of sub-genomic RNA (sgRNA) (50), whereby nested species of sub-genomic messenger RNA (sgmRNA) are produced. Structural proteins of 229E are translated from sgmRNA 2, 4, 5, 6, 7, while sgmRNA 3 is truncated and was described to be non-functional (51–53). The leader-body junction consists of nucleotides within the transcription-regulating sequence of the leader (TRS-L) and the TRS of the body (TRS-B). The hybridization process is mediated by long range RNA-RNA interaction that leads to the formation of sgmRNA which can be distinguished based on their sequence from the genomic RNA. Reverse primers specific for the spike (226R) and ORF4 (617R) genes were designed to synthesize cDNA from total RNA extracted from infected cell lysates. Using a semi-nested PCR approach, the 2F primer targeting the start of the leader RNA was used in combination with 158R, and subsequently with 87R to generate PCR products that contain the leader RNA, leader-body junction, and a portion of sgmRNA 2. Similarly, to test the presence of sgmRNA 4, 2F was employed together with primers 520R and 397R. PCR products were visualized on a 1.5% agarose gel.

### Protein modelling

The full genomic sequences of human coronavirus 229E C2R10 isolates were translated to amino acid sequences using the Expasy Translate webserver (https://web.expasy.org/translate/) (54) in all three forward 5’ to 3’ reading frames. These amino acid sequences were then aligned to the full-length sequences of 229E spike and ORF4 proteins obtained from GenBank (accession number: NC_002645) using Clustal Omega webserver (https://www.ebi.ac.uk/Tools/msa/clustalo/) (55) to determine reading frames corresponding to each protein and deleted residues. The full-length structural models of human coronavirus 229E spike protein was built using the SARS-CoV-2 spike protein model as the template (56). Modeller version 9.21 (57) was used to generate ten homology models and the best model was selected based on having the lowest discreet optimized protein energy (58), while minimizing Ramachandran outliers (59). Using similar protocols, the ORF4 protein model was built using the cryo-EM structure of SARS-CoV-2 ORF3a ion channel (PDB: 7KJR) (60) as the template. The N- and C-terminal regions were unresolved and therefore modelled as unstructured loops. Deleted regions of the C2R10 isolates were mapped to these structures and visualized with VMD (61).

### Polyclonal serum generation and virus neutralization test (VNT)

Recombinant human coronavirus 229E spike protein (Sinobiological, China) was used to immunize two New Zealand rabbits thrice with two-week intervals. Whole blood was subjected to antibody affinity purification. Caco2 and Rhileki cells were seeded at a concentration of 2×10^6^ cells prior to infection. Polyclonal antisera were serially diluted two-fold and were incubated with 126 TCID_50_ of C2, C2R10, and C2R10C1 viruses for 1 hour at 33°C. Subsequently, Caco2 and Rhileki cells were inoculated for 2 hours at 33°C, media was replenished, and virus titers were quantified at 3 dpi.

### Western blot

Supernatants harvested from infected Rhileki cells were centrifuged at 3,500 rpm and precipitated with 10% polyethylene glycol 8000 (Merck, Germany) overnight in 4°C. The solution was centrifuged at 12,000 rpm for 1 hour at 4°C and pellet resuspended in NTE buffer (1M NaCl, 1M Tris-HCl pH 8.0, 0.5M EDTA pH 8.0). The suspension was subjected to a 20% sucrose cushion via ultracentrifugation using a SW41 rotor at 32,000 rpm for 2 hours at 4°C. The pellet was resuspended in NTE buffer and reduced with 5x lane marker sample buffer (ThermoFisher, USA) at 95°C for 10 minutes. Purified virions were separated on a 4 - 20% Mini-PROTEAN TGX stain-free protein gel (Bio-Rad, USA). The gel was transferred on a polyvinylidene fluoride membrane and blocked with 5% BSA at room temperature for 1 hour. This was followed by overnight probing with primary antibody at 4°C and 1 hour with anti-mouse and anti-rabbit horseradish peroxidase-linked IgG secondary antibodies. Blots were visualized using a CCD imager detector (Bio-Rad, USA) after ECL detection (Amersham, United Kingsom). The primary antibodies include anti-229E nucleocapsid mouse monoclonal (Eurofins Ingenasa, Spain) and anti-229E spike rabbit polyclonal antibodies (Genscript, USA).

### Sialidase treatment

Caco2 and Rhileki cells were seeded in 48-well plates one day prior to treatment. On the day of inoculation, cells were washed with PBS once and treated with *Vibrio Cholerae* neuraminidase at concentrations of 1mU, 2.5mU, and 5mU for 3 hours at 37°C.

## Acknowledgments

We acknowledge John A. Crump, Jenny G.H. Low, Venance P. Maro, Eng Eong Ooi, and Matt P. Rubach for contributing patient specimens from which 229E viruses were isolated. We thank Edward C. Holmes and Raoul J. de Groot for helpful comments and discussions.

## Author Contributions

ML and GJDS designed and supervised the research. MGM, ML, DHWL, and ZY conducted experiments and performed genome sequencing. MGM, JJ and FYW performed genome assembly and annotation. MGM, ML, JJ, FYW, IHM, YCFS, and GJDS performed the genome analysis and interpretation. FS and PJB conducted molecular modelling. MGM, ML, and GJDS wrote the paper. All authors took part in data interpretation and edited the manuscript.

## References

1. Gorbalenya AE, Baker SC, Baric RS, De Groot RJ, Drosten C, Gulyaeva AA, et al. The species Severe acute respiratory syndrome-related coronavirus: classifying 2019-nCoV and naming it SARS-CoV-2. Nat. Microbiol. 5, 536–544. Link: https://gonaturecom/3cW9qJR. 2020;

2. Paules CI, Marston HD, Fauci AS. Coronavirus Infections-More Than Just the Common Cold. JAMA. 2020 Feb 25;323(8):707–8.

3. Nickbakhsh S, Ho A, Marques DFP, McMenamin J, Gunson RN, Murcia PR. Epidemiology of Seasonal Coronaviruses: Establishing the Context for the Emergence of Coronavirus Disease 2019. J Infect Dis. 2020 Jun 16;222(1):17–25.

4. Peiris JSM, Lai ST, Poon LLM, Guan Y, Yam LYC, Lim W, et al. Coronavirus as a possible cause of severe acute respiratory syndrome. Lancet. 2003 Apr 19;361(9366):1319–25.

5. Ksiazek TG, Erdman D, Goldsmith CS, Zaki SR, Peret T, Emery S, et al. A novel coronavirus associated with severe acute respiratory syndrome. N Engl J Med. 2003 May 15;348(20):1953–66.

6. Zaki AM, van Boheemen S, Bestebroer TM, Osterhaus ADME, Fouchier RAM. Isolation of a novel coronavirus from a man with pneumonia in Saudi Arabia. N Engl J Med. 2012 Nov 8;367(19):1814–20.

7. Zhu N, Zhang D, Wang W, Li X, Yang B, Song J, et al. A Novel Coronavirus from Patients with Pneumonia in China, 2019. N Engl J Med. 2020 Feb 20;382(8):727–33.

8. Forni D, Cagliani R, Clerici M, Sironi M. Molecular Evolution of Human Coronavirus Genomes. Trends Microbiol. 2017 Jan;25(1):35–48.

9. Cui J, Li F, Shi ZL. Origin and evolution of pathogenic coronaviruses. Nat Rev Microbiol. 2019 Mar;17(3):181–92.

10. Pfefferle S, Oppong S, Drexler JF, Gloza-Rausch F, Ipsen A, Seebens A, et al. Distant Relatives of Severe Acute Respiratory Syndrome Coronavirus and Close Relatives of Human Coronavirus 229E in Bats, Ghana. Emerg Infect Dis. 2009;15(9):1377–84.

11. Montecino-Latorre D, Goldstein T, Gilardi K, Wolking D, Van Wormer E, Kazwala R, et al. Reproduction of East-African bats may guide risk mitigation for coronavirus spillover. One Health Outlook. 2020 Feb 7;2(1):2.

12. Lau SKP, Lung DC, Wong EYM, Aw-Yong KL, Wong ACP, Luk HKH, et al. Molecular Evolution of Human Coronavirus 229E in Hong Kong and a Fatal COVID-19 Case Involving Coinfection with a Novel Human Coronavirus 229E Genogroup. mSphere. 2021 Feb 10;6(1):e00819–20.

13. Crossley BM, Mock RE, Callison SA, Hietala SK. Identification and Characterization of a Novel Alpaca Respiratory Coronavirus Most Closely Related to the Human Coronavirus 229E. Viruses. 2012 Dec;4(12):3689–700.

14. Sabir JSM, Lam TTY, Ahmed MMM, Li L, Shen Y, Abo-Aba SEM, et al. Co-circulation of three camel coronavirus species and recombination of MERS-CoVs in Saudi Arabia. Science. 2016 Jan 1;351(6268):81–4.

15. Vijaykrishna D, Smith GJD, Zhang JX, Peiris JSM, Chen H, Guan Y. Evolutionary insights into the ecology of coronaviruses. J Virol. 2007 Apr;81(8):4012–20.

16. Irving AT, Ahn M, Goh G, Anderson DE, Wang LF. Lessons from the host defences of bats, a unique viral reservoir. Nature. 2021 Jan;589(7842):363–70.

17. Menachery VD, Graham RL, Baric RS. Jumping species—a mechanism for coronavirus persistence and survival. Curr Opin Virol. 2017 Apr;23:1–7.

18. Li W, Moore MJ, Vasilieva N, Sui J, Wong SK, Berne MA, et al. Angiotensin-converting enzyme 2 is a functional receptor for the SARS coronavirus. Nature. 2003 Nov 27;426(6965):450–4.

19. Hoffmann M, Kleine-Weber H, Schroeder S, Krüger N, Herrler T, Erichsen S, et al. SARS-CoV-2 Cell Entry Depends on ACE2 and TMPRSS2 and Is Blocked by a Clinically Proven Protease Inhibitor. Cell. 2020 Apr 16;181(2):271–280.e8.

20. Hofmann H, Pyrc K, van der Hoek L, Geier M, Berkhout B, Pöhlmann S. Human coronavirus NL63 employs the severe acute respiratory syndrome coronavirus receptor for cellular entry. Proc Natl Acad Sci U S A. 2005 May 31;102(22):7988–93.

21. Raj VS, Mou H, Smits SL, Dekkers DHW, Müller MA, Dijkman R, et al. Dipeptidyl peptidase 4 is a functional receptor for the emerging human coronavirus-EMC. Nature. 2013 Mar 14;495(7440):251–4.

22. Yeager CL, Ashmun RA, Williams RK, Cardellichio CB, Shapiro LH, Look AT, et al. Human aminopeptidase N is a receptor for human coronavirus 229E. Nature. 1992 Jun 4;357(6377):420–2.

23. Zhang R, Wang K, Lv W, Yu W, Xie S, Xu K, et al. The ORF4a protein of human coronavirus 229E functions as a viroporin that regulates viral production. Biochim Biophys Acta. 2014 Apr;1838(4):1088–95.

24. Koch J, Uckeley ZM, Doldan P, Stanifer M, Boulant S, Lozach PY. TMPRSS2 expression dictates the entry route used by SARS-CoV-2 to infect host cells. EMBO J. 2021 Aug 16;40(16):e107821.

25. Liu K, Pan X, Li L, Yu F, Zheng A, Du P, et al. Binding and molecular basis of the bat coronavirus RaTG13 virus to ACE2 in humans and other species. Cell. 2021 Jun 24;184(13):3438–3451.e10.

26. Wells HL, Letko M, Lasso G, Ssebide B, Nziza J, Byarugaba DK, et al. The evolutionary history of ACE2 usage within the coronavirus subgenus Sarbecovirus. Virus Evol. 2021 Jan;7(1):veab007.

27. Starr TN, Zepeda SK, Walls AC, Greaney AJ, Alkhovsky S, Veesler D, et al. ACE2 binding is an ancestral and evolvable trait of sarbecoviruses. Nature. 2022 Feb 3;1–9.

28. Hoffmann M, Müller MA, Drexler JF, Glende J, Erdt M, Gützkow T, et al. Differential Sensitivity of Bat Cells to Infection by Enveloped RNA Viruses: Coronaviruses, Paramyxoviruses, Filoviruses, and Influenza Viruses. PLoS One. 2013 Aug 30;8(8):e72942.

29. Letko M, Marzi A, Munster V. Functional assessment of cell entry and receptor usage for SARS-CoV-2 and other lineage B betacoronaviruses. Nat Microbiol. 2020 Apr;5(4):562–9.

30. Ren W, Qu X, Li W, Han Z, Yu M, Zhou P, et al. Difference in receptor usage between severe acute respiratory syndrome (SARS) coronavirus and SARS-like coronavirus of bat origin. J Virol. 2008 Feb;82(4):1899–907.

31. Khaledian E, Ulusan S, Erickson J, Fawcett S, Letko MC, Broschat SL. Sequence determinants of human-cell entry identified in ACE2-independent bat sarbecoviruses: A combined laboratory and computational network science approach. eBioMedicine. 2022 May 1;79:103990.

32. Gallagher TM, Buchmeier MJ, Perlman S. Cell receptor-independent infection by a neurotropic murine coronavirus. Virology. 1992 Nov;191(1):517–22.

33. Menachery VD, Dinnon KH, Yount BL, McAnarney ET, Gralinski LE, Hale A, et al. Trypsin Treatment Unlocks Barrier for Zoonotic Bat Coronavirus Infection. J Virol. 2020 Feb 14;94(5):e01774–19.

34. Shen XR, Geng R, Li Q, Chen Y, Li SF, Wang Q, et al. ACE2-independent infection of T lymphocytes by SARS-CoV-2. Sig Transduct Target Ther. 2022 Mar 11;7(1):1–11.

35. Xiong Q, Cao L, Ma C, Liu C, Si J, Liu P, et al. Close relatives of MERS-CoV in bats use ACE2 as their functional receptors [Internet]. bioRxiv; 2022 [cited 2022 May 27]. p. 2022.01.24.477490. Available from: https://www.biorxiv.org/content/10.1101/2022.01.24.477490v1

36. Munster VJ, Adney DR, van Doremalen N, Brown VR, Miazgowicz KL, Milne-Price S, et al. Replication and shedding of MERS-CoV in Jamaican fruit bats (Artibeus jamaicensis). Sci Rep. 2016 Feb 22;6(1):21878.

37. Morandi MI, Busko P, Ozer-Partuk E, Khan S, Zarfati G, Elbaz-Alon Y, et al. Extracellular vesicle fusion visualized by cryo-EM [Internet]. bioRxiv; 2022 [cited 2022 May 27]. p. 2022.03.28.486013. Available from: https://www.biorxiv.org/content/10.1101/2022.03.28.486013v1

38. Freeman MC, Peek CT, Becker MM, Smith EC, Denison MR. Coronaviruses induce entry-independent, continuous macropinocytosis. mBio. 2014 Aug 5;5(4):e01340–01314.

39. Lim JP, Gleeson PA. Macropinocytosis: an endocytic pathway for internalising large gulps. Immunol Cell Biol. 2011 Nov;89(8):836–43.

40. Plowright RK, Eby P, Hudson PJ, Smith IL, Westcott D, Bryden WL, et al. Ecological dynamics of emerging bat virus spillover. Proc Biol Sci. 2015 Jan 7;282(1798):20142124.

41. Goh G, Ahn M, Zhu F, Lee LB, Luo D, Irving AT, et al. Complementary regulation of caspase-1 and IL-1β reveals additional mechanisms of dampened inflammation in bats. Proc Natl Acad Sci U S A. 2020 Nov 17;117(46):28939–49.

42. Latinne A, Hu B, Olival KJ, Zhu G, Zhang L, Li H, et al. Origin and cross-species transmission of bat coronaviruses in China. Nat Commun. 2020 Aug 25;11(1):4235.

43. Low JGH, Ooi EE, Tolfvenstam T, Leo YS, Hibberd ML, Ng LC, et al. Early Dengue infection and outcome study (EDEN) - study design and preliminary findings. Ann Acad Med Singap. 2006 Nov;35(11):783–9.

44. Reed LJ, Muench H. A Simple Method of Estimating Fifty Per Cent Endpoints. American Journal of Epidemiology. 1938 May 1;27(3):493–7.

45. Bolger AM, Lohse M, Usadel B. Trimmomatic: a flexible trimmer for Illumina sequence data. Bioinformatics. 2014 Aug 1;30(15):2114–20.

46. Okonechnikov K, Golosova O, Fursov M, Ugene team. Unipro UGENE: a unified bioinformatics toolkit. Bioinformatics. 2012 Apr 15;28(8):1166–7.

47. Robinson JT, Thorvaldsdóttir H, Winckler W, Guttman M, Lander ES, Getz G, et al. Integrative Genomics Viewer. Nat Biotechnol. 2011 Jan;29(1):24–6.

48. Katoh K, Standley DM. MAFFT Multiple Sequence Alignment Software Version 7: Improvements in Performance and Usability. Molecular Biology and Evolution. 2013 Apr 1;30(4):772–80.

49. Katoh K, Misawa K, Kuma K ichi, Miyata T. MAFFT: a novel method for rapid multiple sequence alignment based on fast Fourier transform. Nucleic Acids Res. 2002 Jul 15;30(14):3059–66.

50. Pasternak AO, Spaan WJM, Snijder EJ. Nidovirus transcription: how to make sense…? J Gen Virol. 2006 Jun;87(Pt 6):1403–21.

51. Viehweger A, Krautwurst S, Lamkiewicz K, Madhugiri R, Ziebuhr J, Hölzer M, et al. Direct RNA nanopore sequencing of full-length coronavirus genomes provides novel insights into structural variants and enables modification analysis. Genome Res. 2019 Sep;29(9):1545–54.

52. Schreiber SS, Kamahora T, Lai MM. Sequence analysis of the nucleocapsid protein gene of human coronavirus 229E. Virology. 1989 Mar;169(1):142–51.

53. Raabe T, Schelle-Prinz B, Siddell SG. Nucleotide sequence of the gene encoding the spike glycoprotein of human coronavirus HCV 229E. J Gen Virol. 1990 May;71 (Pt 5):1065–73.

54. Gasteiger E, Gattiker A, Hoogland C, Ivanyi I, Appel RD, Bairoch A. ExPASy: The proteomics server for in-depth protein knowledge and analysis. Nucleic Acids Res. 2003 Jul 1;31(13):3784–8.

55. Sievers F, Wilm A, Dineen D, Gibson TJ, Karplus K, Li W, et al. Fast, scalable generation of high-quality protein multiple sequence alignments using Clustal Omega. Mol Syst Biol. 2011 Oct 11;7:539.

56. Raghuvamsi PV, Tulsian NK, Samsudin F, Qian X, Purushotorman K, Yue G, et al. SARS-CoV-2 S protein:ACE2 interaction reveals novel allosteric targets. Elife. 2021 Feb 8;10:e63646.

57. Sali A, Blundell TL. Comparative protein modelling by satisfaction of spatial restraints. J Mol Biol. 1993 Dec 5;234(3):779–815.

58. Eramian D, Shen M yi, Devos D, Melo F, Sali A, Marti-Renom MA. A composite score for predicting errors in protein structure models. Protein Sci. 2006 Jul;15(7):1653–66.

59. Ramachandran GN, Ramakrishnan C, Sasisekharan V. Stereochemistry of polypeptide chain configurations. J Mol Biol. 1963 Jul;7:95–9.

60. Kern DM, Sorum B, Mali SS, Hoel CM, Sridharan S, Remis JP, et al. Cryo-EM structure of SARS-CoV-2 ORF3a in lipid nanodiscs. Nat Struct Mol Biol. 2021 Jul;28(7):573–82.

61. Humphrey W, Dalke A, Schulten K. VMD: visual molecular dynamics. J Mol Graph. 1996 Feb;14(1):33–8, 27–8.

