## Supplementary online material for "Spike-independent infection of human coronavirus 229E in bat cells"

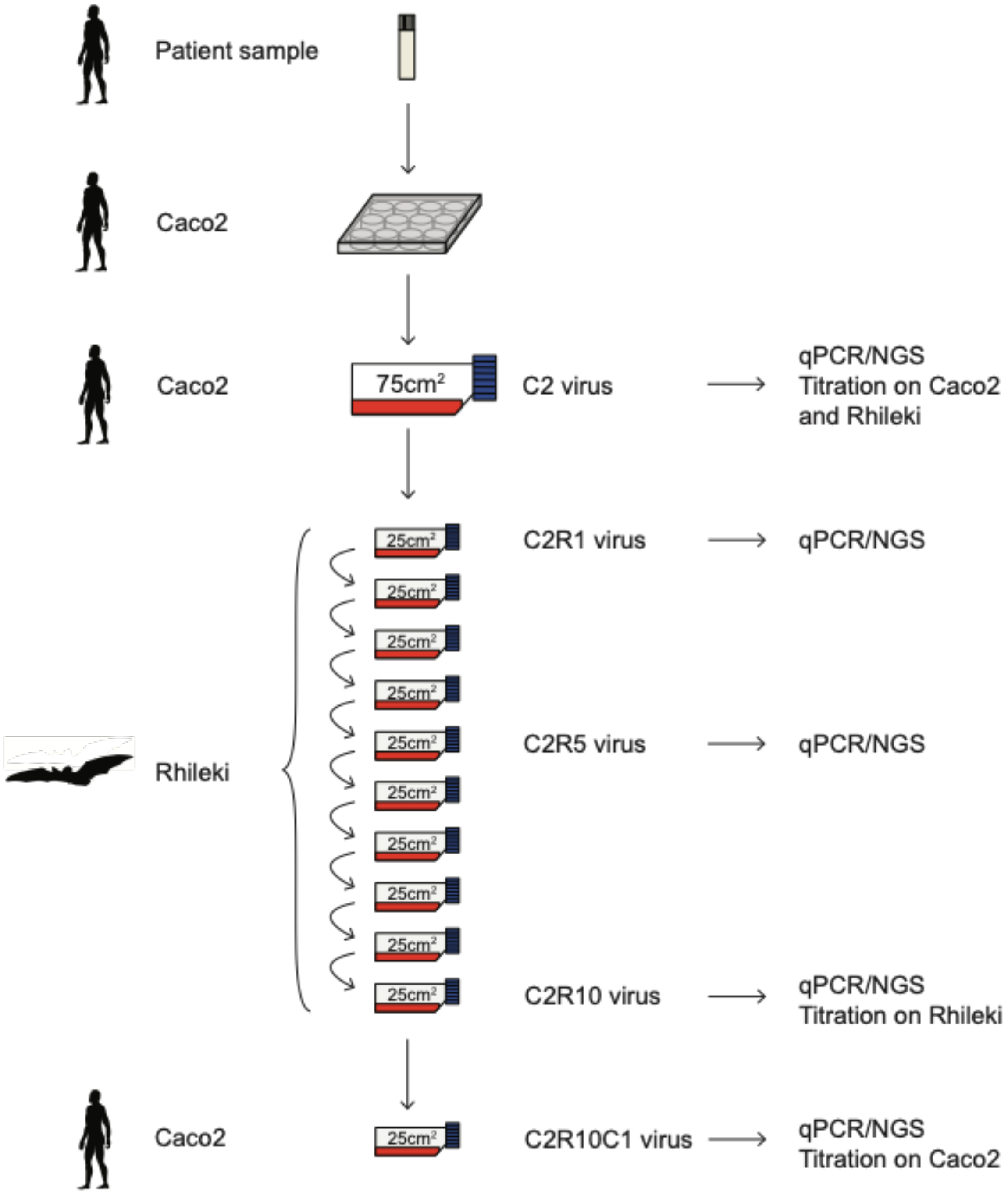


**Figure S1. Schematic of virus passage experiments.** Succession of the virus cultures, nomenclature of viruses, and tests performed on individual isolates.

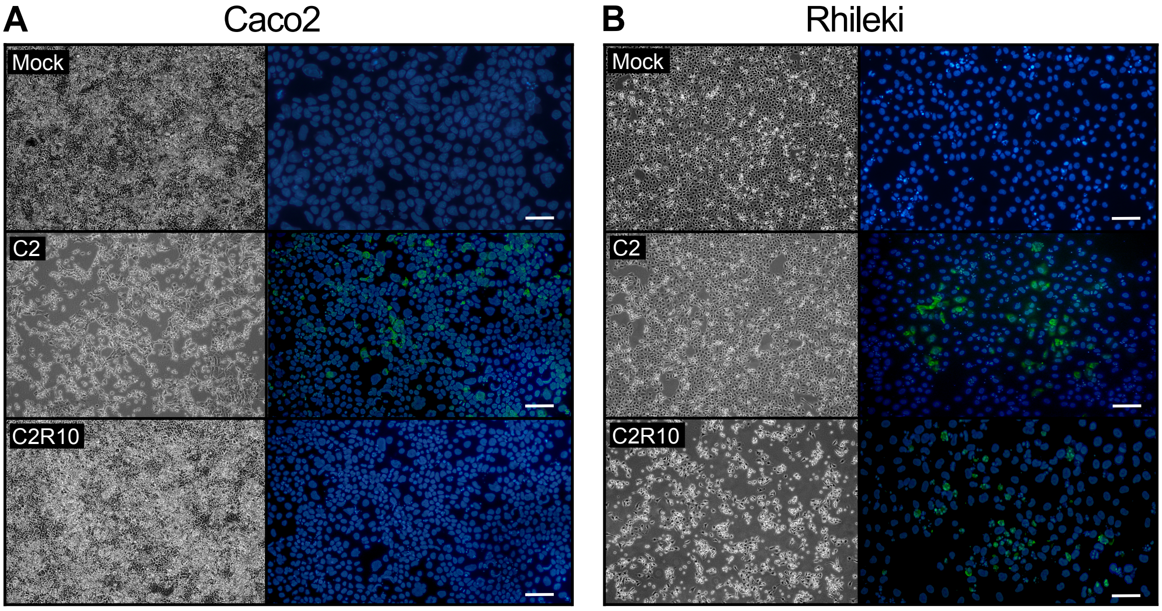


**Figure S2. Microscopy and immunofluorescence of Caco2 and Rhileki cells infected with C2 or C2R10 viruses.** (A) Bright light and immunofluorescence microscopy of Caco2 cells inoculated with C2, C2R10 virus or mock. (B) Bright light and immunofluorescence microscopy of Rhileki cells inoculated with C2, C2R10 virus or mock. 229E viral nucleocapsid staining (green) and nuclear staining (blue). White scale bars correspond to 100 µm.

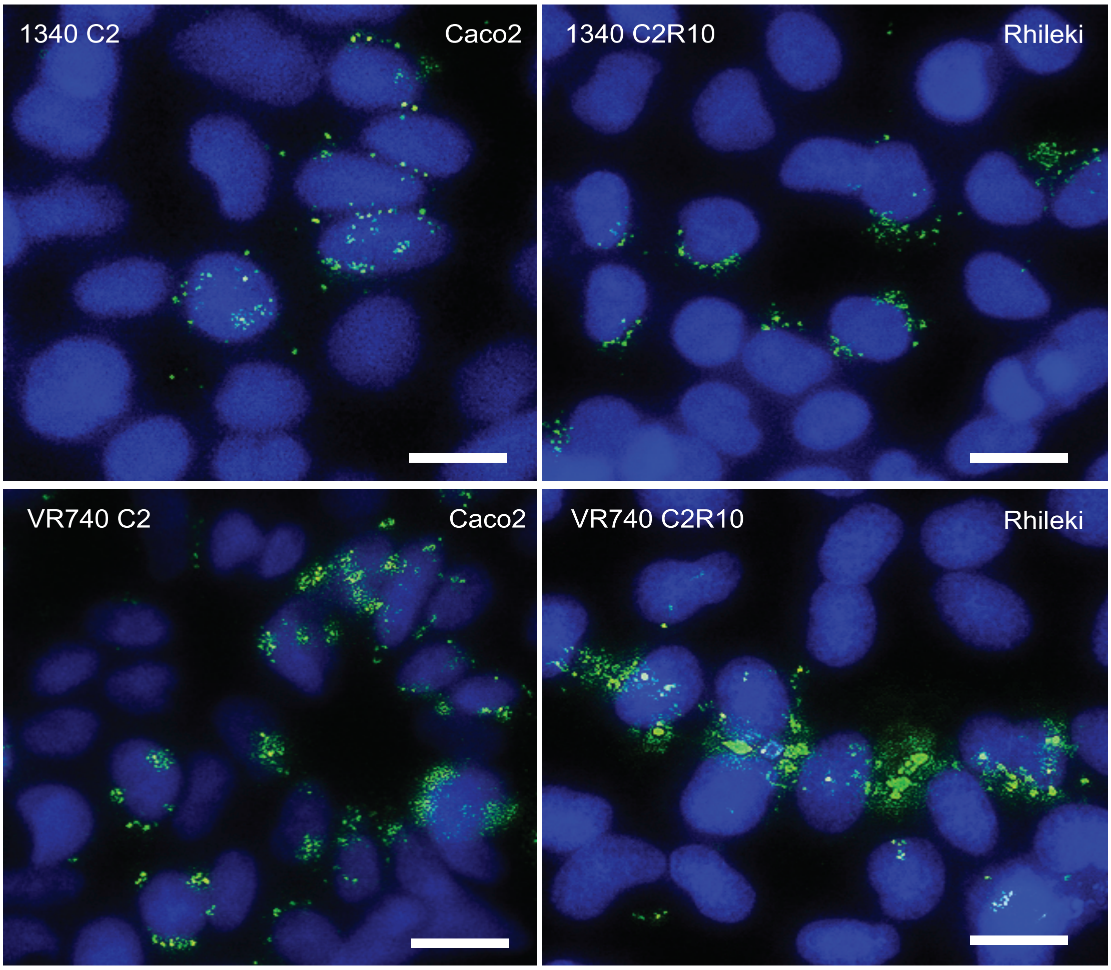


**Figure S3. Double-stranded RNA staining of Caco2 and Rhileki cells.** Representative images of Caco2 cells inoculated with selected C2 viruses and Rhileki cells inoculated with two C2R10 viruses. Double-stranded RNA intermediates (green) and nuclear staining (blue) visualized at 1 day post inoculation. White scale bars correspond to 20 µm.

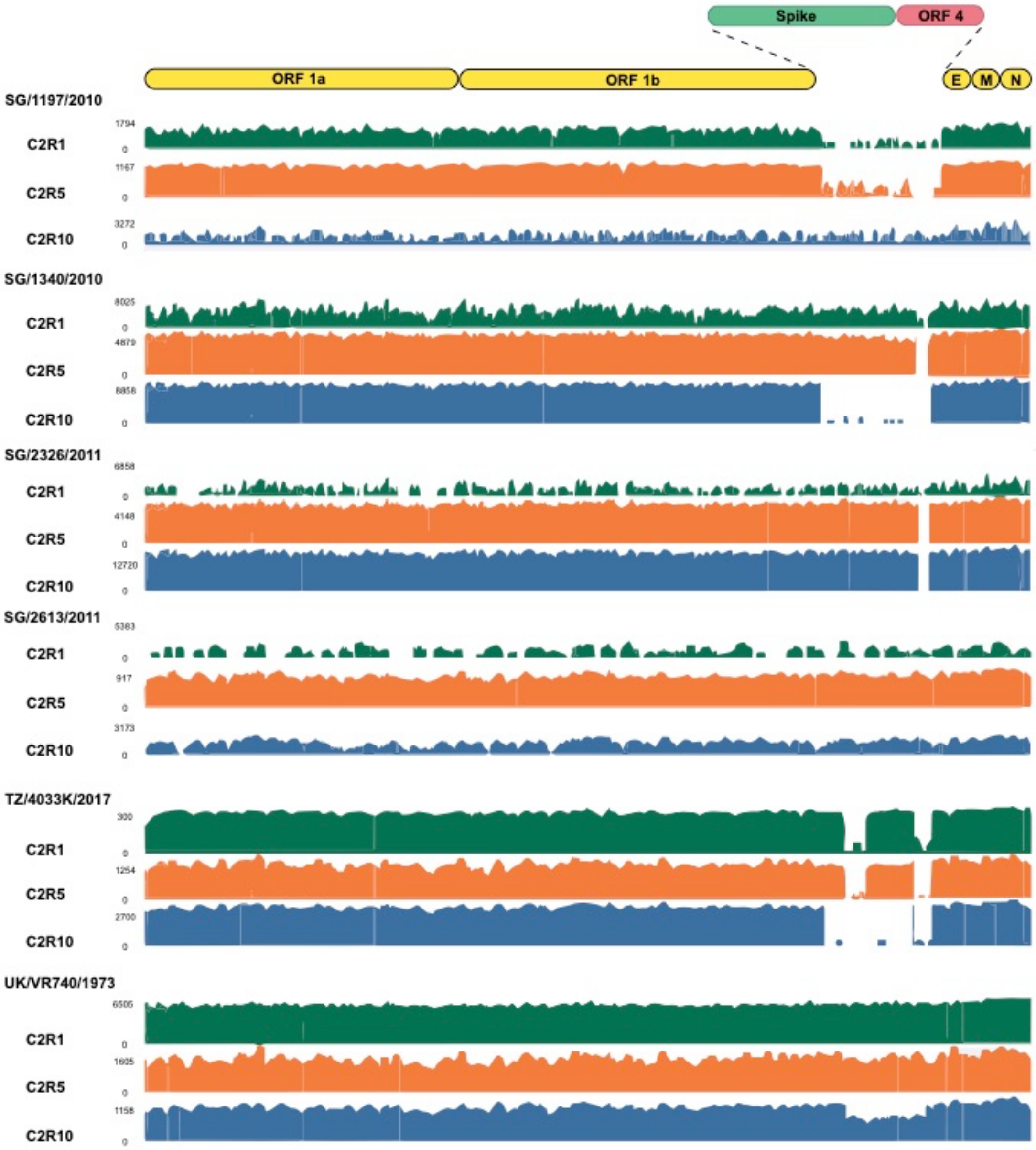


**Figure S4. Full genome coverage plot of next-generation sequencing reads obtained from Rhileki cells infected with C2R1, C2R5, or C2R10 viruses.**

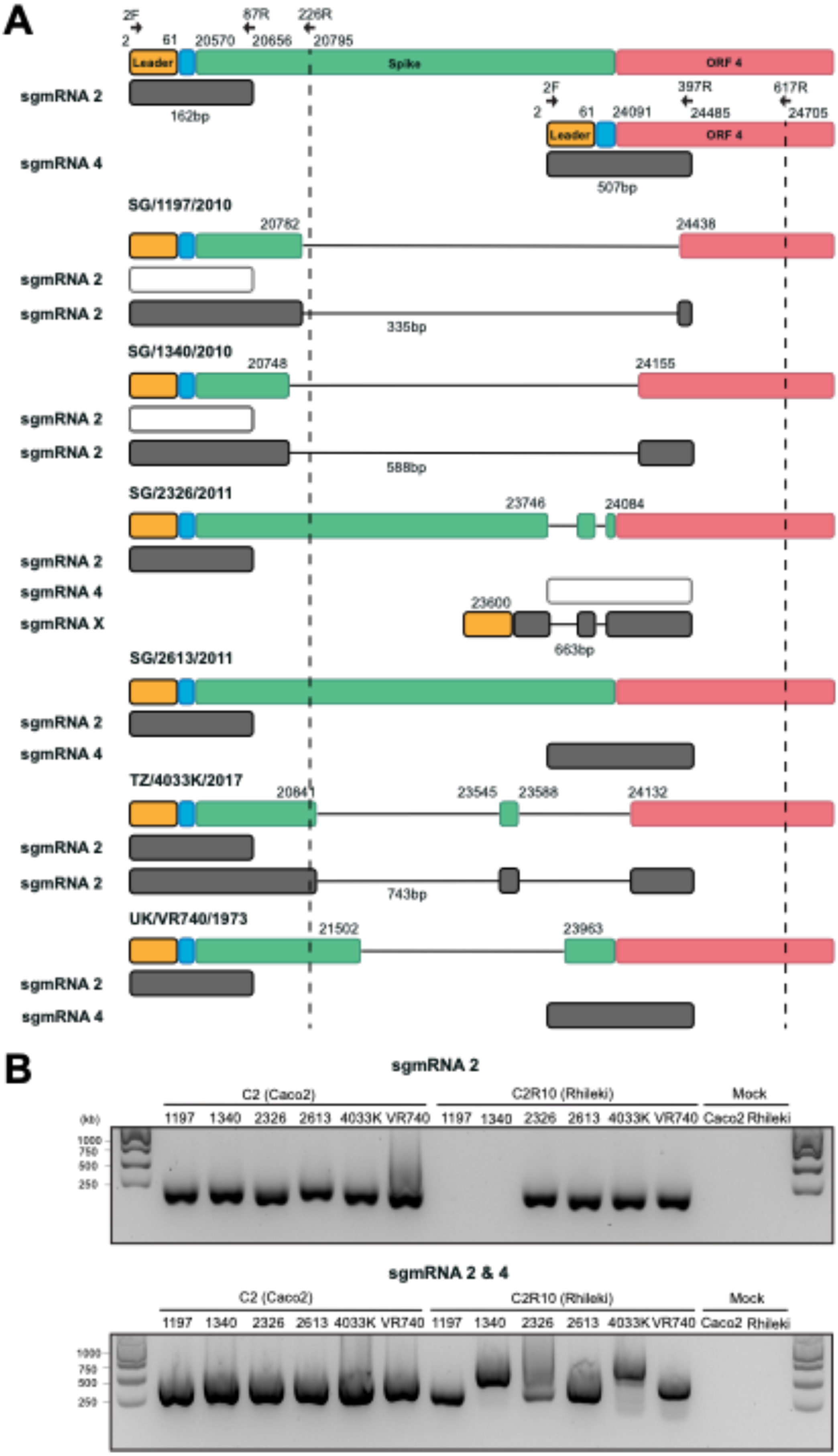
A
2F
87R
226R
2
61
20570
20656
20795
Leader
Spike
ORF 4
sgmRNA 2
162bp
2F
397R
617R
2
61
24091
24485
24705
Leader
ORF 4
sgmRNA 4
507bp
SG/1197/2010
20782
24438
sgmRNA 2
sgmRNA 2
335bp
SG/1340/2010
20748
24155
sgmRNA 2
sgmRNA 2
588bp
SG/2326/2011
23746
24084
sgmRNA 2
sgmRNA 4
sgmRNA X
23600
663bp
SG/2613/2011
sgmRNA 2
sgmRNA 4
TZ/4033K/2017
20841
23545
23588
24132
sgmRNA 2
sgmRNA 2
743bp
UK/VR740/1973
21502
23963
sgmRNA 2
sgmRNA 4
B
sgmRNA 2
C2 (Caco2)
C2R10 (Rhileki)
Mock
(kb)
1197 1340 2326 2613 4033K VR740 1197 1340 2326 2613 4033K VR740 Caco2 Rhileki
1000
750
500
250
sgmRNA 2 & 4
C2 (Caco2)
C2R10 (Rhileki)
Mock
1197 1340 2326 2613 4033K VR740 1197 1340 2326 2613 4033K VR740 Caco2 Rhileki
1000
750
500
250

**Figure S5. Schematic describing the detection of 229E subgenomic mRNA 2 and 4 in total RNA extracted from C2R10 virus infected Rhileki cells.** (A) Filled rectangles represent nucleotides at the 5’ end of sgmRNA 2 and 4 species. The leader sequence, leader-body junction (LBJ), spike, and ORF 4 reading frames are represented in orange, blue, green, and red, respectively. Primers targeting the LBJ of both sgmRNA 2 and 4 are denoted by arrows. PCR products are indicated by dark gray rectangles for sgmRNA 2 (162bp) and sgmRNA 4 (507bp), whereas white rectangles represent the absence of an expected PCR product. Rectangles of varying sizes represent the observed sequences in spike and ORF4 regions transcribed from genomic RNA. Horizontal black lines indicate deletions. (B) PCR amplification and gel electrophoresis of PCR products described in (A). Total RNA extracted Caco2 cells infected with C2 viruses were used for comparison purposes.

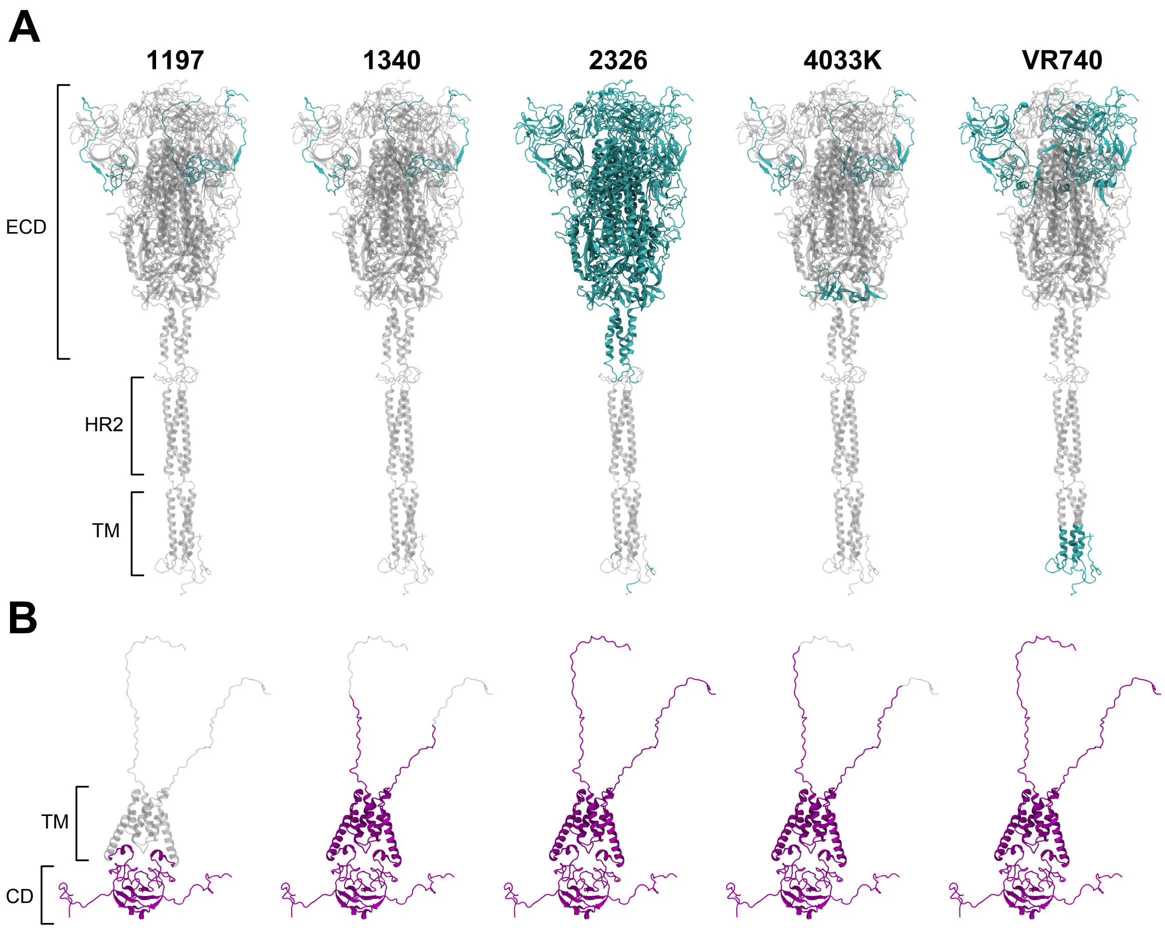


**Figure S6. Modelling of deleted regions of the spike and ORF4 proteins of C2R10**. (A) Full-length model of 229E spike protein built using SARS-CoV-2 spike protein as a structural template. The model is shown in cartoon representation, with the deleted regions coloured in grey, and remaining spike protein in cyan. (B) Full-length model of ORF4 protein, built using the cryo-EM structure of the SARS-CoV-2 ORF3a ion channel (PDB: 7KJR). Deleted regions are shown in grey, and the remaining protein shown in purple. Domains are labelled as ECD (ectodomain), HR2 (heptad repeat 2 domain), TM (transmembrane domain), and CD (cytoplasmic domain).

**Table S1. Primers used in this study**.

| Name | Directionality | Sequence | Usage |
|---|---|---|---|
| 1197 P5 Del F | Forward | CGGCTGTGTTGGTCATTCG | Deletion verification |
| 1197 P5 Del R | Reverse | GGCTGAGGTCTTTATCAATACG | Deletion verification |
| 1340 P5 Del F | Forward | CGTATTCCTACCATAGC | Deletion verification |
| 1340 P5 Del R | Reverse | AGATCTGCCACGGTGTGATC | Deletion verification |
| 2326 P5 Del F | Forward | GCTCTGAGTTGCAAACTATTGTGC | Deletion verification |
| 2326 P5 Del R | Reverse | CGTGAAACTTCAGCACTAAC | Deletion verification |
| T140 P5 Del F | Forward | CTTGTTAGGAGTGGTAAGTTGC | Deletion verification |
| T140 P5 Del R | Reverse | GCAAGTTGAAGGTAACAGTGCC | Deletion verification |
| T140 P10 Del F | Forward | GCATATGCCTTGTTGCATATTGC | Deletion verification |
| T140 P10 Del R | Reverse | CGCTAGCAAGTTGAAGGTAACAGTGC | Deletion verification |
| T140 Del R | Reverse | GCACAACCCAGACCACG | Deletion verification |
| T4033 P5 Del F | Forward | CTCCATACTGTCTTGCTGC | Deletion verification |
| T4033 P5 Del R | Reverse | GCAGATCTGCCACGGTGTGAT | Deletion verification |
| T4033 P10 Del F | Forward | GGTTATATACCCTCCAACTTTGC | Deletion verification |
| T4033 P1-1 Del F | Forward | AGACTGAGATGTGACCAGC | Deletion verification |
| T4033 P1-2 Del F | Forward | TTCGTGCTTCCAGACAGCTTGC | Deletion verification |
| T4033 P1-1 Del R | Reverse | GCATGGCGCTATTTCTTAAGGC | Deletion verification |
| T4033 P10 Del-2 F | Forward | CGCATAATGTTTGAACCACG | Deletion verification |
| 2 F | Forward | CTTAAGTACCTTATCTATCTACAGATA | sgmRNA 2/4 PCR 1/2 |
| 226 R | Reverse | GCTGAAAACTCCTCACAACACC | sgmRNA 2 cDNA |
| 158 R | Reverse | GCAAAGTTGGAGGGTATATAACC | sgmRNA 2 PCR 1 |
| 87 R | Reverse | GCAAACAGAGTGACTAGTGTTCGTC | sgmRNA 2 PCR 2 |
| 617 R | Reverse | GCTGGGTGTTCACAAAACAAGTATA | sgmRNA 4 cDNA |
| 520 R | Reverse | CGTACAAATCGTTAGTTGAGAG | sgmRNA 4 PCR 1 |
| 397 R | Reverse | GCACATAGCAAAGTGTGGTTAC | sgmRNA 4 PCR 2 |

**Table S2. Assembly and statistics of NGS reads including reads mapping in deleted areas.**

| | | SG/1197/2010 | SG/1340/2010 | SG/2326/2011 | TZ/4033K/2017 | UK/VR740/1973 |
|---|---|---|---|---|---|---|
| Deletion present? | | Y | N | N | Y | N |
| Size of deletion (nt) | | 2878 | - | - | 449 | - |
| # of C2R1 NGS reads | C2 virus consensus | 348 | 1114 | 102 | 12836 | 29608 |
| mapping to | C2R1 consensus | 344 | 1114 | 102 | 12701 | 29608 |
| | Deleted region | 4/348 (1.15%) | NA | NA | 135/12836 (1.05%) | NA |
| Ratio of reads relative to size of deletion | | $3.9x10^{-6}$ | NA | NA | $3.9x10^{-5}$ | NA |
| Deletion present? | | Y | Y | Y | Y | N |
| Size of deletion (nt) | | 3656 | 443 | 324 | 1207 | - |
| # of C2R5 NGS reads | C2 virus consensus | 2897 | 48069 | 44157 | 22486 | 30839 |
| mapping to | C2R5 consensus | 2877 | 48862 | 43840 | 22175 | 30839 |
| | Deleted region | 20/2897 (0.69%) | 207/49069 (0.65%) | 317/44157 (0.72%) | 311/22486 (1.38%) | NA |
| Ratio of reads relative to size of deletion | | $1.9x10^{-6}$ | $9.5x10^{-6}$ | $2.2x10^{-5}$ | $1.1x10^{-5}$ | NA |
| Deletion present? | | Y | Y | Y | Y | Y |
| Size of deletion (nt) | | 3656 | 3407 | 324 | 3248 | 2461 |
| # of C2R10 NGS reads | C2 virus consensus | ND | 64463 | 51642 | 70566 | 17096 |
| mapping to | C2R10 consensus | ND | 63981 | 51205 | 63250 | 16644 |
| | Deleted region | ND | 482/64464 (0.74%) | 437/51642 (0.85%) | 1316/70566 (1.86%) | 452/17096 (2.64%) |
| Ratio of reads relative to size of deletion | | ND | $2.2x10^{-6}$ | $2.6x10^{-5}$ | $5.7x10^{-6}$ | $1.1x10^{-5}$ |

*ND – Not done due to low coverage

*NA – Not applicable

**Table S3. Synonymous and non-synonymous mutations in virus from passages 1, 5, and 10.**

| Isolate | Gene | Nucleotide mutation | Amino acid mutation | Mutation | C2 | C2R1 | C2R5 | C2R10 |
|---|---|---|---|---|---|---|---|---|
| SG/1340/2010 | NSP3 | A4854G | D621G | Non-synonymous | D | 100% D | 65.5% G | 100% G |
| | NSP10 | C12249T | T46I | Non-synonymous | T | 47% T | 64.0% I | 100% I |
| SG/2326/2011 | NSP3 | A4854G | D621G | Non-synonymous | D | 100% D | 100% D | 100% G |
| | NSP15 | T19179G | - | Synonymous | T | 100% G | 95.0% G | 100% G |
| | Membrane | C25177T | L54F | Non-synonymous | L | 100% L | 100% F | 100% F |
| | Membrane | T25258C | F81L | Non-synonymous | F | 100% F | 100% F | 100% L |
| SG/2613/2011 | NSP8 | C11462T | H88Y | Non-synonymous | H | 100% Y | 100% Y | 100% Y |
| | Spike | C20603T | L5F | Non-synonymous | L | 100% L | 100% F | 100% F |
| | ORF4 | C24200T | Q30* | Non-synonymous | Q | 100% * | 100% * | 100% * |
| | ORF4 | G24423T | G104V | Non-synonymous | G | 100% G | 96.0% G | 100% V |
| TZ/4033K/2017 | NSP2 | C2954T | - | Synonymous | C | 100% T | 100% T | 100% T |
| | NSP3 | C7074T | S1361F | Non-synonymous | S | 100% F | 100% F | 100% F |
| | Membrane | C25177T | L54F | Non-synonymous | L | 100% L | 100% L | 100% F |
| | Nucleocapsid | T26118C | V137A | Non-synonymous | V | 100% V | 97.1% V | 100% A |
| UK/VR740/1973 | NSP2 | C712T | - | Synonymous | C | 100% T | 72.4% T | 100% T |
| | NSP2 | C1083A | S153Y | Non-synonymous | S | 100% S | 100% S | 86.9% Y |
| | NSP3 | A4907G | I634V | Non-synonymous | I | 62.05% I | 100% V | 100% V |
| | NSP4 | T7848C | M28T | Non-synonymous | M | 51.7% T | 100% T | 100% T |

*SG/1197/2010 was excluded due to low coverage
